# Unravelling the complex genetic basis of growth in New Zealand silver trevally (*Pseudocaranx georgianus*)

**DOI:** 10.1101/2021.10.11.463933

**Authors:** Noemie Valenza-Troubat, Sara Montanari, Peter Ritchie, Maren Wellenreuther

## Abstract

Growth directly influences production rate and therefore is one of the most important and well-studied trait in animal breeding. However, understanding the genetic basis of growth has been hindered by its typically complex polygenic architecture. Here, we performed quantitative trait locus (QTL) mapping and genome-wide association studies (GWAS) for 10 growth traits that were observed over two years in 1,100 F_1_ captive-bred trevally (*Pseudocaranx georgianus*). We constructed the first high-density linkage map for trevally, which included 19,861 single nucleotide polymorphism (SNP) markers, and discovered eight QTLs for height, length and weight on linkage groups 3, 14 and 18. Using GWAS, we further identified 113 SNP-trait associations, uncovering 10 genetic hot spots involved in growth. Two of the markers found in the GWAS co-located with the QTLs previously mentioned, demonstrating that combining QTL mapping and GWAS represents a powerful approach for the identification and validation of loci controlling complex traits. This is the first study of its kind for trevally. Our findings provide important insights into the genetic architecture of growth in this species and supply a basis for fine mapping QTLs, marker-assisted selection, and further detailed functional analysis of the genes underlying growth in trevally.

## Introduction

Growth facilitates essential functions such as reproduction or the ability to adapt to environments. Although there are exceptions, increase in body size is usually positively correlated with numerous fitness traits such as higher mating success and fecundity, increased offspring quality and lengthened longevity (Gebhardt-Henrich and Richner 1998, Dmitriew 2011). Most vertebrates exhibit a finite amount of growth. In fish, however, the process of growing is indeterminate, or indefinite, and continues throughout their life (Weatherley 1972), although the rate tends to decline as body size increases (Pedersen and Jobling 1989). This highly complex process is the result of interactions between environmental effects and genetic differences. Factors including sex (e.g. Imsland and Jonassen 2003), age or food availability (e.g. Jones 1986) influence growth rate, as well as abiotic factors changing seasonally such as temperature (e.g. Karås and Klingsheim 1997, Imsland, Schram et al. 2007), photoperiod (e.g. Imsland and Jonassen 2003) or oxygen levels (e.g. Brett and Groves 1979). The genetic basis of growth traits is typically highly polygenic (Wellenreuther and Hansson 2016). Functional relationships between genetic variations and physiological parameters of growth have been described in commercially relevant species such as cod (*Gadus morhua*) (see review by: Imsland and Jónsdóttir 2002), Atlantic salmon (*Salmo salar*) (Tsai, Hamilton et al. 2015) or tilapia (*Tilapia mossambica*) (Liu, Sun et al. 2014), but are still poorly understood in many non-model species.

With the development of next generation sequencing technologies (NGS) the cost of high-density genotyping has drastically decreased, therefore enabling the more widespread application of quantitative trait locus (QTL) mapping experiments. Many variants associated with complex phenotypes have since been found in a variety of non-model species. However, QTL mapping approaches only allow the identification of genetic regions that are polymorphic between two parents, and relevant in a particular environment, therefore potentially missing some common variants associated with the traits (Mackay 2001). For this reason the location and effects of detected QTLs can vary between mapping populations. For example, in the Atlantic salmon, QTLs involved in growth have been found on different linkage groups according to different studies (Baranski, Moen et al. 2010, Tsai, Hamilton et al. 2015, Besnier, Solberg et al. 2020).

Recently, genome-wide association studies (GWAS) have been widely employed to detect QTLs both in captive (Palaiokostas, Kocour et al. 2018) and natural populations (Santure and Garant 2018). This technique identifies associations between markers and phenotypes based on linkage disequilibrium (LD) (Meuwissen and Goddard 2010). Compared with QTL mapping, GWAS can be conducted on genetically diverse, unrelated individuals and is particularly advantageous when controlled crossing and generation of large segregating populations is difficult. GWAS has proved useful for the identification of loci associated with numerous growth traits in fish (e.g. Yang, Wu et al. 2020). However, the power of GWAS relies on the number of markers in relation to the extent of linkage disequilibrium in the population (Newell, Cook et al. 2011). When applied to natural populations or collections of outcrossing individuals, characterised by a rapid linkage disequilibrium decay, tens of thousands of markers are often required to obtain an adequate level of resolution. Because of the intrinsic limitations of statistical methods comparing thousands of tests, GWAS has a higher frequency of false discovery of loci than QTL mapping (Hayes, Lewin et al. 2013). A final disadvantage of GWAS is its low power for detecting rare allelic variants. However, their frequency can be increased in controlled crosses and therefore captured with QTL mapping. Hence, combining GWAS and QTL mapping often gives more complete and reliable results than using one of the methods alone (Fraslin, Quillet et al. 2020).

Aquaculture is the fastest growing food-production sector in Aotearoa New Zealand. Currently, the local industry relies almost exclusively on the farming of three species: Greenshell™ mussels (*Perna canaliculus*), Pacific oysters (*Crassostrea gigas*) and only one finfish, king/chinook salmon (*Oncorhynchus tshawytscha*), which is an introduced species (Camara and Symonds 2014, Symonds, Clarke et al. 2019). Hence, there is a strong interest in diversifying the range of species available for aquaculture. The native finfish silver trevally (*Pseudocaranx georgianus*, Cuvier 1833) has been identified as a suitable candidate for New Zealand aquaculture. In Aotearoa, its Māori name is araara. Indigenous Māori people have a strong cultural connection to trevally, where it is considered as taonga (i.e. has value, or is treasured). Trevally is a shoaling pelagic species found throughout the coastal waters of southern Australia and around New Zealand (Smith-Vaniz and Jelks 2006). It is most common at depths of approximately 80 m, although its range is thought to be 10–238 m and can reach over 40 years of age (Ministry for Primary Industries 2014, Bray 2020). In many regions of New Zealand, trevally is a major component of recreational and commercial fisheries (Ministry for Primary Industries 2014). For it to be suitable for commercial aquaculture, however, trevally’s growth rate must be improved, which can be done using selective breeding (Valenza-Troubat, Montanari et al. 2021).

In this study, we investigated the genetic architecture of 10 growth traits in a new population of trevally. More specifically, we genotyped and phenotyped 1,100 F_1_ trevally to 1) generate a high-density linkage map, 2) detect rare QTLs associated with growth using linkage mapping in a sub-family comprising 89 individuals, 3) identify common SNPs associated with growth using GWAS on the whole population, and 4) overlap the results and annotate supported regions.

## Methods

### Study population

The trevally population used in this study was generated as part of a breeding programme started at the Plant and Food Research (PFR) finfish research facility in Nelson, New Zealand. The full description of holding conditions and pedigree are described in Valenza-Troubat *et al*. (2021). Briefly, the population comprised of 13 wild caught F_0_ broodstock and 1,100 F_1_ captive-bred offspring. In 2015, induced mass spawning of the F_0_ generation in a single tank was used to produce the offspring F_1_ generation. This resulted in a complex pedigree, including a combination of unrelated, full- and half-siblings in the F_1_ generation. F_1_ offspring were held in a single tank from hatch receiving the same feeding regime, light, water flow and aeration until the end of this experiment. The seawater tanks in the facility are located on the seaward side of Port Nelson and receive ambient seawater from an underground bore which is filtered using mesh filters and UV treatment.

### Phenotyping and trait estimation

Ten growth traits were used in the current study, namely peduncle length (PL), height at 25 (H25), 50 (H50) and 75% (H75) of the PL, estimated weight (EW) and related net gain traits (ΔPL, ΔH25, ΔH50, ΔH75 and ΔEW, respectively) (Valenza-Troubat, Montanari et al. 2021). These measurements were recorded on three occasions throughout the experiment, at the beginning (November 2017), in the middle (October 2018), and at the end (November 2019), when the fish were a little over two, three and four years old, respectively. Using the Morphometric Software™ (https://www.plantandfood.co.nz/page/morphometric-software-home/), the outline of each fish was extracted from images and morphometric were located, and then used to make measurements. PL was measured by assessing the distance between the upper lip and narrowest cross-section of the tail. Height was measured at three positions along the fish: 25, 50, and 75% of the PL. The weight estimations (EW) were done following a Bayesian hierarchical approach as described in Froese et al. (2014). The net gain for each trait was calculated as the difference from the initial measurement (November 2017) with subsequent ones. The normality of the data was assessed visually using Quantile-Quantile (QQ) plots generated in the R statistical environment version 3.2.3 (R Development Core Team 2016).

### Genotyping and variant calling

Samples of fin tissue from 13 tagged F_0_ and 1,100 F_1_ individuals were collected and stored as described in Valenza-Troubat et al. (2021). Total DNA was extracted as described in Ashton et al. (2019) with minor modifications, and then quantified and quality-checked by fluorescence, spectrophotometry and agarose gel electrophoresis. The 13 F_0_ individuals were whole genome sequenced (paired-end, 125 bp reads) over three lanes of the HiSeq 2500 platform at the Australian Genome Research Facility (AGRF, Melbourne, Australia). The F_1_ were genotyped using a modified GBS approach (Elshire, Glaubitz et al. 2011), as described in Valenza-Troubat *et al*. (2021). A total of 12 pools of 96 samples each were prepared and sent to AGRF for sequencing on a HiSeq 2500 platform (single-end, 100 bp reads). Sequencing data quality for both F_0_ and F_1_ generations were checked using FastQC v0.11.7 (Andrews 2010). Raw reads from the F_0_ were trimmed using trimmomatic v0.36 (Bolger, Lohse et al. 2014) (using the parameters HEADCROP: 9, TRAILING: 10, SLIDINGWINDOW: 5:20, MINLEN: 75). The F_1_ samples were de-multiplexed from the 12 sequencing libraries using the process_radtags module available in the STACKs version 2.1 pipeline (Catchen, Hohenlohe et al. 2013) and the reads were trimmed using Fastq-mcf in ea-utils v1.1.2-806 (minimum sequence length = 50, quality threshold causing base removal = 33) (Aronesty 2013). Read groups were added to all sequences and bam files were sorted and indexed using Picard toolkit (Broad_Institute 2015). Reads were then mapped to the reference genome (Catanach, Ruigrok et al. 2021) using the Burrows-Wheeler Aligner (BWA) v0.7.17 (Li & Durbin, 2009) and variants were called jointly with the parallel module of freebayes v1.3.1 (Garrison and Marth 2012), with minimum of five observations and minimum mapping quality of 10. The SNPs with single-sample sequence coverage (sequencing depth) < 3 were removed to reduce the number of putatively erroneous genetic variants, and missing data and minor allele frequency (MAF) were set to < 20% and > 0.05, respectively.

### Linkage map construction and QTL identification

The parents of each F_1_ individual in the dataset were identified with Sequoia v2.0.7 (Huisman 2017) as reported in Valenza-Troubat *et al*. (2021). The full SNP dataset (i.e. before the stringent filtering performed for the parentage analysis) was filtered for Mendelian errors (> 5%), and checked for distorted segregation using a chi-square test with α=0.05. The linkage map was constructed in Lep-MAP v3.0 (Rastas, Paulin et al. 2013), using the largest family. Markers were separated into linkage groups (LG) with the SeparateChromosomes module (logarithm of odds (LOD) limit = 14, minimum markers per LG = 50). The order of the markers was computed with the OrderMarkers module. Single markers at the end of each LG were removed if they were more than 3 cM apart from the next closest marker. MapChart v2.32 (Voorrips 2002) was used to visualise the genetic map. As the F_0_ were assumed to be outbred, the linkage map and the genotypes of the mapping family were input as a 4-way cross in the R package R/qtl version 1.47-9 (Broman, Wu et al. 2003) for interval mapping. Standard interval mapping was performed and a genome-wide permutation test (Doerge and Churchill 1996) with 1000 permutations was used to determine the LOD significance thresholds (*p*-value = 0.05).

### Genome-Wide Association Study

GWAS was carried out on the entire genotyped F_1_ population (n=1100). SNPs were removed if the call rate was smaller than 0.8, MAF < 0.01, if Mendel error rate > 5% (based on trios identified with the parentage analysis carried out above), and they were LD-pruned using an r^2^ > 0.80 in a 50kb sliding window with five variants. Association analysis were performed using the Fixed and random model Circulating Probability Unification (FarmCPU) method (Liu, Huang et al. 2016) implemented in GAPIT3 v3.1.0 (Wang and Zhang 2021). A Bayesian information criterion (BIC)-based model selection was used to find the optimal number of principal components (PCs) for each time measure, to account for family and population structure. The cut-off for significant association was a False Discovery Rate (FDR) adjusted *p*-value = 0.05 (Benjamini and Hochberg 1995), to control for multiple testing. To assess how well the model used in GWAS accounted for population structure and family relatedness, results of the GWAS were visualised with QQ plots implemented in GAPIT3, which depicted the distribution of the actual *p*-values compared with the theoretical ones. Manhattan plots were used to visualise the SNPs associated with the different phenotypes, using the physical position of the markers on the reference genome.

### Ethics statement

All research carried out in this study was reviewed and approved by the animal ethics committee of Victoria University of Wellington in New Zealand (Application number 25976).

## Results

### Phenotypic values of growth traits

All offspring were phenotyped at the first measurement (November 2017), while numbers decreased at the two subsequent time points because of natural mortality that occurred during the study. All traits showed a normal distribution and exhibited large levels of phenotypic variation, as discussed in Valenza-Troubat et al. (2021) A summary of the mean values and standard deviations of the 10 growth phenotypes is shown in Table 1. In both the family used for QTL mapping and in the entire population used for GWAS, a normal distribution of the residuals was observed for the ten traits investigated (Figure 1). Transgressive lines (i.e. offspring that have more extreme phenotypes than the parents) were observed in the population.

**Table 1:** Summary of the phenotypes height at 25%, 50% and 75% of the body (H25, H50 and H75 respectively), Peduncle Length (PL), estimated weight (EW) and net gains traits associated (ΔH25, ΔH50, ΔH74, ΔPL and ΔEW respectively) across the whole F1 population and for the largest family. Included are the number offspring phenotyped (n), mean, and standard deviation (SD).

|  | Nov 2017 |  | Oct 2018 |  | Nov 2019 |  |
| --- | --- | --- | --- | --- | --- | --- |
| *All* | n = 1,094 |  | n= 719 |  | n= 694 |  |
|  | Mean | SD | Mean | SD | Mean | SD |
| H25 | 49.71 | 9.02 | 66.94 | 13.51 | 90.10 | 15.70 |
| H50 | 59.45 | 11.02 | 79.99 | 16.39 | 105.53 | 18.65 |
| H75 | 48.58 | 9.91 | 61.26 | 12.69 | 85.63 | 15.80 |
| PL | 156.79 | 23.92 | 200.64 | 35.36 | 264.09 | 40.44 |
| EW | 89.90 | 41.85 | 191.93 | 99.89 | 426.34 | 177.93 |
| ΔH25 | | | 15.85 | 7.80 | 39.08 | 11.21 |
| ΔH50 | | | 18.75 | 8.42 | 44.34 | 12.36 |
| ΔH75 | | | 11.08 | 6.77 | 35.54 | 10.19 |
| ΔPL | | | 40.03 | 18.00 | 103.59 | 27.31 |
| ΔEW | | | 95.83 | 64.45 | 330.78 | 147.12 |
| *Family* | n = 87 |  | n = 68 |  | n = 67 |  |
|  | Mean | SD | Mean | SD | Mean | SD |
| H25 | 52.21 | 7.95 | 73.71 | 11.32 | 96.52 | 12.64 |
| H50 | 63.10 | 9.29 | 86.87 | 13.41 | 111.90 | 14.77 |
| H75 | 52.85 | 8.99 | 66.72 | 9.75 | 92.36 | 12.39 |
| PL | 167.59 | 21.32 | 218.19 | 30.04 | 280.78 | 33.07 |
| EW | 107.43 | 37.85 | 238.01 | 89.61 | 498.52 | 159.82 |
| ΔH25 | | | 20.43 | 6.53 | 42.78 | 9.21 |
| ΔH50 | | | 22.46 | 6.92 | 46.84 | 9.89 |
| ΔH75 | | | 12.88 | 5.54 | 37.79 | 8.33 |
| ΔPL | | | 47.91 | 15.91 | 109.31 | 23.15 |
| ΔEW | | | 125.99 | 60.54 | 384.63 | 135.00 |

**Figure 1:**
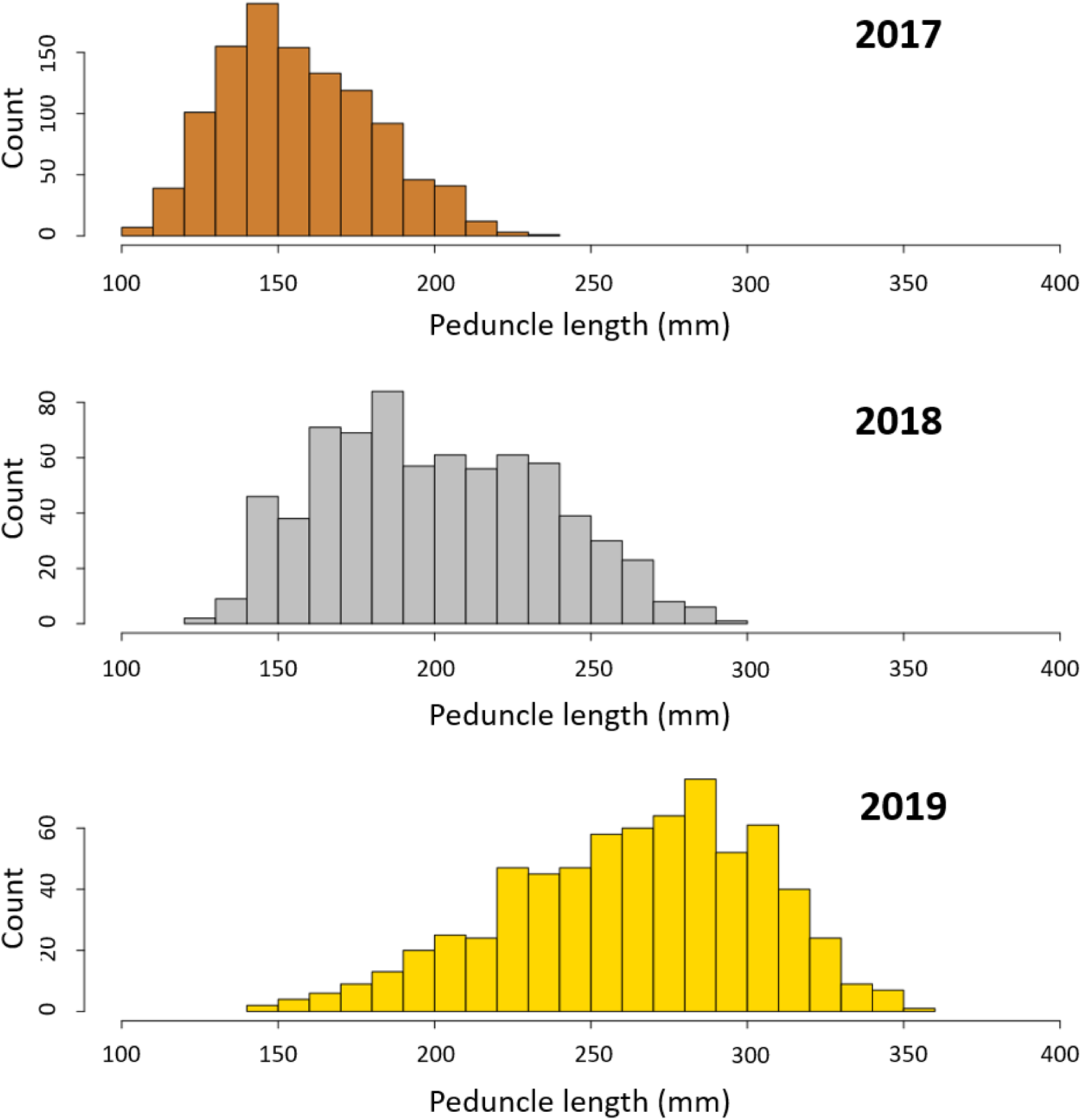
Distribution of the peduncle length measurements in November 2017 (n = 1,094), October 2018 (n= 719) and November 2019 (n = 694).

### High-density linkage map

Sequencing of the 13 parental fishes generated 1.23 billion reads, corresponding to an average genome coverage of 13x per individual. Of the 1,100 offspring, 1094 F1 were successfully genotyped via GBS, resulting in 0.42x genome coverage per individual fish. Variant calling and basic quality filtering generated a dataset of 171,923 SNPs. Overall, 21 families from 10 out of the 13 sequenced F0 fishes were identified by Valenza-Troubat et al. (2021) when reconstructing the pedigree of the population. The largest family included 87 offspring and was used to assemble the sex-averaged linkage map. A total of 21,665 SNPs were polymorphic in this family and passed the chi-square test, and 19,861 were successfully mapped to 24 Linkage groups, 1,830 after removal of the co-mapping loci (Figure 2). LG numbers were randomly assigned, as no previous reference had been published. The genetic map spanned 1,335.46 cM, with an average marker distance of 0.73 cM. The largest gap was on LG14 and was 4.10 cM long. The longest and shorter Linkage groups were 17 (82.89 cM) and 8 (48.14 cM), respectively (Table 2).

**Table 2:** Marker statistics of the linkage map constructed from the largest trevally family.

| LG | Number of markers<sup>a</sup> | Length (cM) | Average distance between markers (cM) |
| --- | --- | --- | --- |
| 1 | 85 | 61.27 | 0.72 |
| 2 | 80 | 58.41 | 0.73 |
| 3 | 81 | 57.26 | 0.71 |
| 4 | 68 | 49.34 | 0.73 |
| 5 | 81 | 56.69 | 0.70 |
| 6 | 65 | 51.72 | 0.80 |
| 7 | 87 | 60.09 | 0.69 |
| 8 | 70 | 48.14 | 0.69 |
| 9 | 78 | 54.43 | 0.70 |
| 10 | 84 | 57.22 | 0.68 |
| 11 | 82 | 61.88 | 0.75 |
| 12 | 71 | 49.30 | 0.69 |
| 13 | 79 | 59.00 | 0.75 |
| 14 | 64 | 53.51 | 0.84 |
| 15 | 67 | 52.84 | 0.79 |
| 16 | 67 | 56.27 | 0.84 |
| 17 | 108 | 82.89 | 0.77 |
| 18 | 82 | 54.95 | 0.67 |
| 19 | 80 | 53.82 | 0.67 |
| 20 | 71 | 55.04 | 0.78 |
| 21 | 65 | 52.83 | 0.81 |
| 22 | 73 | 49.29 | 0.68 |
| 23 | 63 | 45.39 | 0.72 |
| 24 | 79 | 53.88 | 0.68 |
| **Total** | **1,830** | **1.335.46** | **0.73** |
| <sup>a</sup> count after removal of co-mapping |  |  |  |

**Figure 2:**
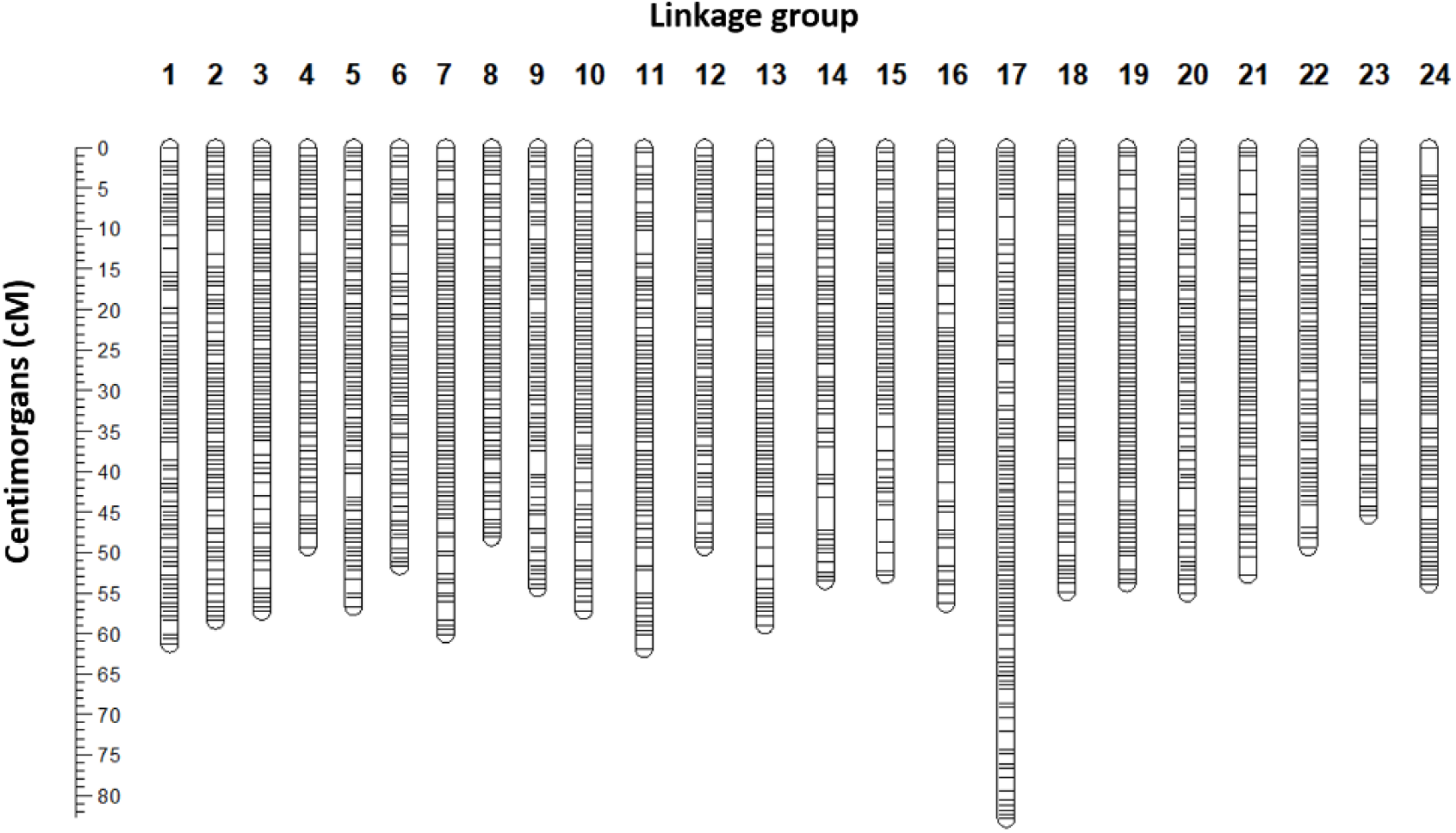
Visualisation of the sex-averaged map built for the largest segregating family (n = 87). The 24 linkage groups represent the expected 24 Pseudocaranx georgianus chromosomes

### QTL mapping revealed rare variants associated with growth

The genome-wide significant thresholds for QTL mapping were established at LOD values between 4.60 and 4.81 for the ten phenotypic traits across three years (Table 3). In total, eight significant QTLs were identified on four locations in three linkage groups (Figure 3). Six of these QTLs were for traits recorded during the first measurement (November 2017), and two for the last measurement (November 2019). In particular, three QTLs detected for H25, PL and EW measured at the first time point co-mapped on LG3 (between 23.2 and 24.9 cM). A second QTL was mapped to LG3 at 52.1 cM for H75 from the first measurement. Additionally, the traits PL and EW from 2017 were associated to a second locus, at the top of LG18. The percentage of phenotypic variance explained (PVE) by the 2017 QTLs ranged from 22.1 to 25.3%. In the last measurement two QTLs for ΔH75 and ΔPL were found to co-map to LG14 (between 17 and 17.1 cM), with PVE of 39.7 and 31.1, respectively (Table 3). No QTLs were found for the middle measurement (October 2018) and for all other traits in 2017 and 2019.

**Table 3:** List of significant Quantitative Trait Loci (QTL) detected for height at 25% and 75% of the peduncle length (H25 and H75 respectively), Peduncle Length (PL), Estimated Weight (EW) and net gain for H75 (ΔH75) and PL (ΔPL) in November 2017 (Nov17) and November 2019 (Nov19), using a high-density linkage map. For each QTL the position on the linkage group (LG), the significance threshold, the LOD at the peak, the SNP name and the percentage of variance explained (PVE) are shown.

| Time | Trait | LG | Peak position (cM) | SNP at peak | LOD threshold | LOD at peak | PVE (%) |
| --- | --- | --- | --- | --- | --- | --- | --- |
| Nov17 | H25 | 3 | 23.2 | trevally000114_4454829 | 4.71 | 4.73 | 22.1 |
| Nov17 | PL | 3 | 24.9 | trevally000114_4498123 | 4.71 | 4.83 | 22.6 |
| Nov17 | EW | 3 | 24.9 | trevally000114_4498123 | 4.60 | 5.39 | 24.8 |
| Nov17 | H75 | 3 | 52.1 | trevally000114_23355470 | 4.64 | 5.50 | 25.3 |
| Nov17 | PL | 18 | 0.0 | trevally001200_2242455 | 4.71 | 4.88 | 22.8 |
| Nov17 | EW | 18 | 0.0 | trevally001200_2242455 | 4.60 | 4.82 | 22.5 |
| Nov19 | ΔPL | 14 | 17.0 | trevally001025_25205374 | 4.81 | 5.41 | 31.0 |
| Nov19 | ΔH75 | 14 | 17.6 | trevally001025_25180484 | 4.69 | 7.36 | 39.7 |

**Figure 3:**
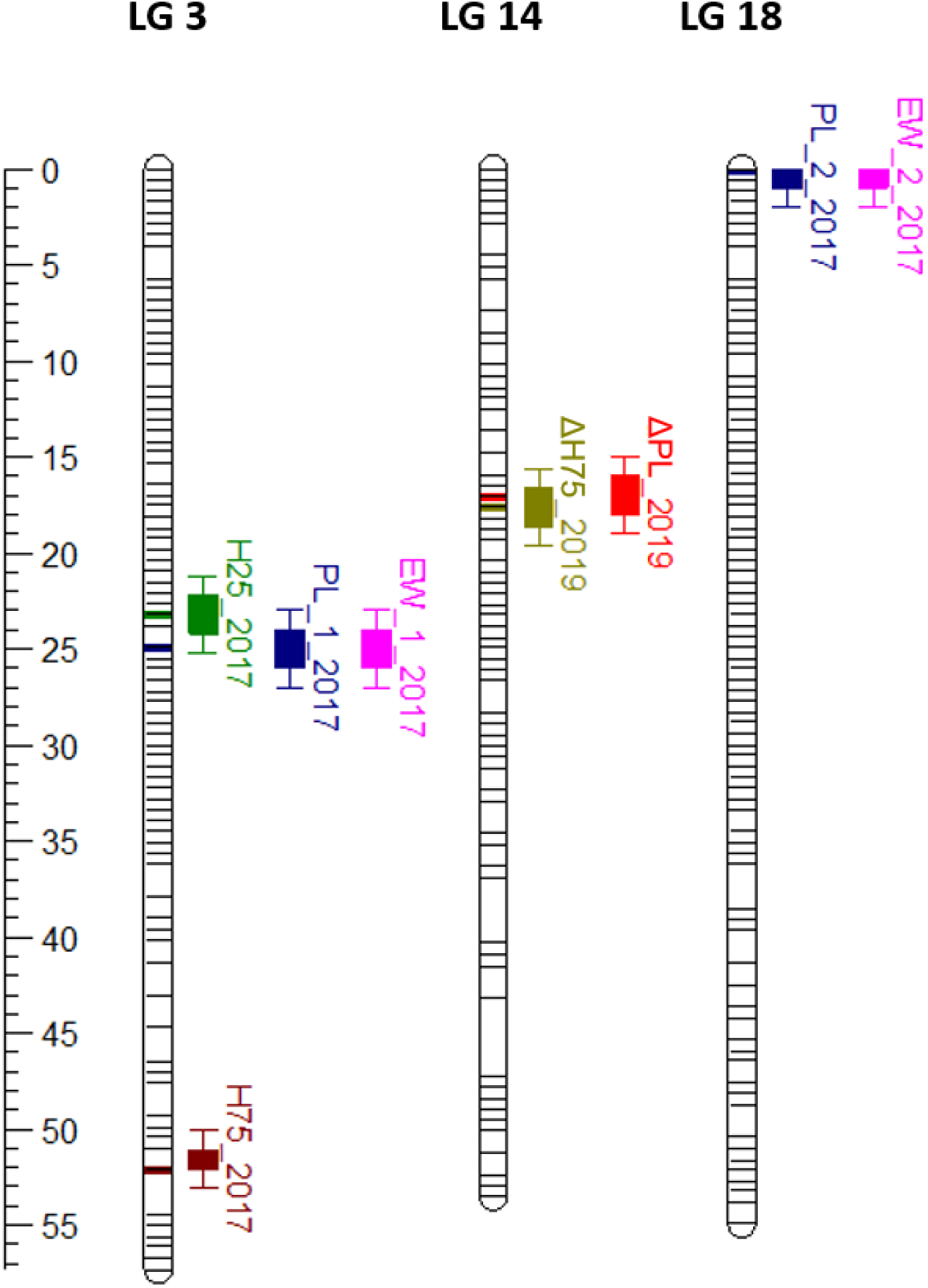
Eight significant Quantitative Trait Loci (QTL) associated with growth traits were found in trevally, across three linkage groups (LG3, LG14 and LG18). In November 2017, two QTLs were found associated with peduncle length (PL_1_2017 and PL_2_2017, on LG3 and LG18 respectively), which were also associated with estimated weight (EW_1_2017 and EW_2_2017). In the same year, two more QTLs were found associated with height at 25% (H75_2017) and 75% (H75_2017) of the peduncle length, both on LG3. Finally, two QTLs were found in November 2019, one for net gain in peduncle length (ΔPL_2019) and one for net gain in height at 75% of the peduncle length (ΔH75_2019), both on LG14.

### Genome-wide association found strongly associated SNPs

A total of 1,024 F1 individuals had both genotypic and phenotypic data for all measurements. After filtering based on Mendelian errors, MAF and LD pruning, 107,067 SNPs were left and used in the GWAS. Model selection in GAPIT resulted in no PCs to be used as co-variates in any of the traits (Figure 4A). QQ plots showed that FarmCPU adequately accounted for the confounding effects of family and population structure (Figure 4B). A total of 93 SNPs were significantly associated (-log10(ρ) > 7.03) with at least one of the ten traits measured at each year of phenotyping (Figure 4C, Table S1). Only ΔH75 in October 2018 and November 2019 had no significant association. Of the 93 SNPs, 15 were associated with more than one trait. These included four, seven and four SNPs associated with measurements at the first, second and third time point, respectively. No common SNPs were identified among different years. Significant associations were in some cases found for SNPs less than 0.5 Mb apart, highlighting hot spots on chromosomic regions. A total of 10 hot spots were identified: two on LG1, one on LG2, two on LG3, two on LG5, one on LG6, one on LG10 and one on LG18. Six of these included associations for traits measured in different years (Table S1).

**Figure 4:**
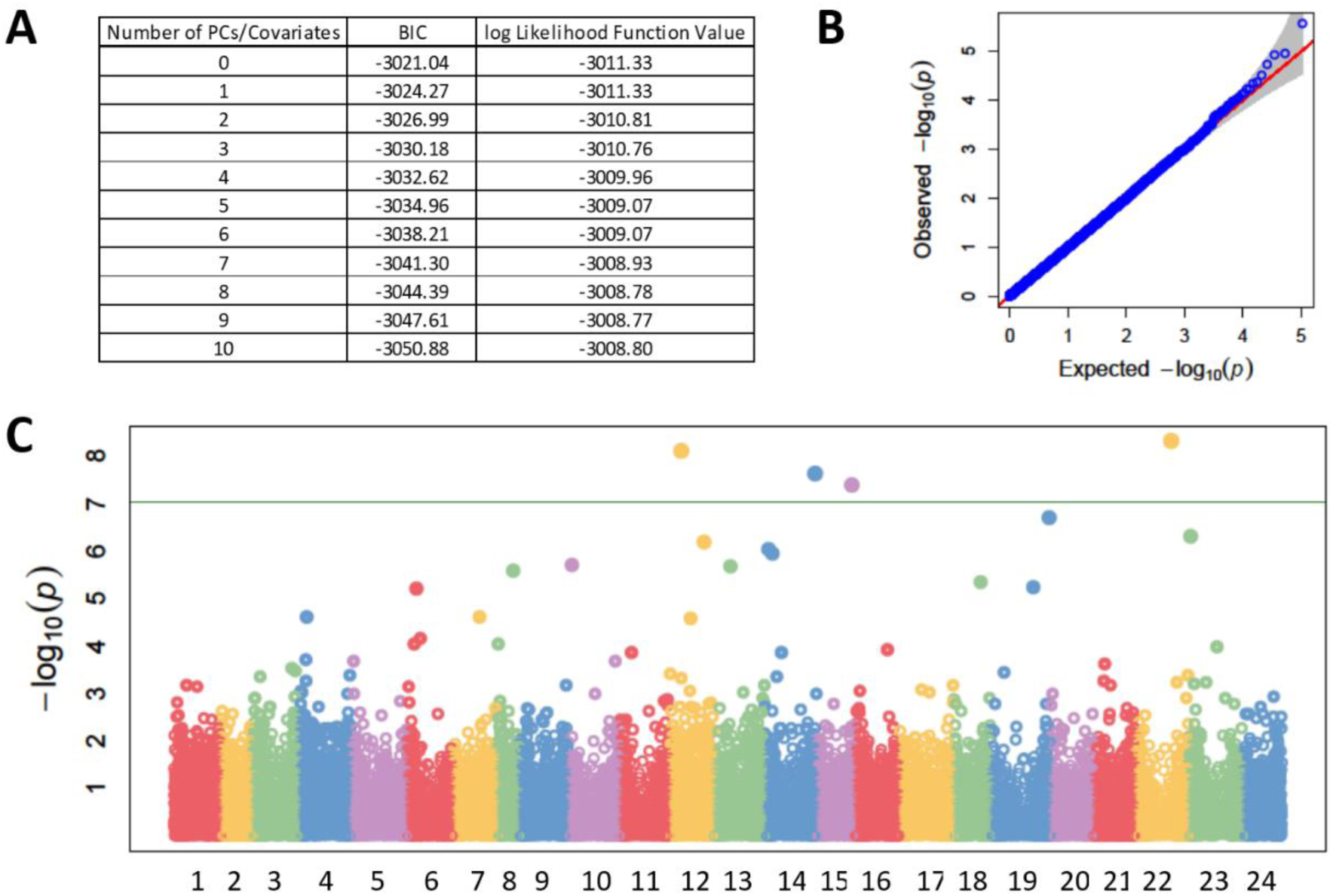
GWAS results for the net gain in peduncle length (ΔPL) in November 2019. (A) Summary of Bayesian information criterion (BIC) for the optimal number of Principal Components (PCs) to use in the model. (B) Quantile-Quantile (QQ)-plot of p-values. On the Y-axis is the observed negative base 10 logarithm of the p-values, and on the X-axis the expected observed negative base 10 logarithm of the p-values under the assumption that the p-values follow a uniform [0 to 1] distribution. The grey area shows the 95% confidence interval for the QQ-plot under the null hypothesis of no association between the SNP and the trait. (C) Manhattan plot of the results of the GWAS. On the X-axis is the physical position of the SNPs on the genome divided by chromosomes, and on the Y-axis is the negative log base 10 of the p-values. The horizontal green line represents the significance threshold.

## Discussion

We constructed the first genetic map for trevally, and we were able to use this map to determine genomic regions associated with phenotypic growth traits. Our map included 19,861 SNPs across 24 linkage groups and confirmed the 24 mega-scaffolds of the trevally reference genome (Catanach, Ruigrok et al. 2021). A linkage map built for yellowtail kingfish (*Seriola lalandi*), the closest species to trevally for which a map has been constructed, was also resolved into 24 linkage groups (Nguyen, Rastas et al. 2018), consistent with our findings. The map built here for trevally was 1,335.46 cM in length, which is within the range of map length expected for many teleost fish. For example, the map assembled here is longer when compared to the yellowtail kingfish map, similar in length to the map of the Australasian snapper (*Chrysophrys auratus*) (Ashton, Ritchie et al. 2019) and shorter when compared to the European sea bass (*Dicentrarchus labrax*) (Griot, Allal et al. 2021).

Eight significant QTLs related to growth were mapped to three of the trevally linkage groups. Other studies in teleost have found multiple QTLs affecting growth such as in Atlantic salmon (Tsai, Hamilton et al. 2015, Besnier, Solberg et al. 2020), Australasian snapper (Ashton, Ritchie et al. 2019), spotted sea bass (*Lateolabrax maculatus*) (Liu, Wang et al. 2020) or yellowtail kingfish (Nguyen, Rastas et al. 2018), supporting the hypothesis of a polygenic regulation. No significant QTLs were identified with QTL mapping for the second time point, October 2018. In the segregating family used for QTL mapping, the number of offspring decreased from 87 in 2017 to 68 in 2018 and 67 in 2019 as the result of natural mortality. Although some statistically significant QTLs were found in the measurements made in the latter period of the experiments, the smaller sample sizes in the two last measurements reduced the power of the study, which could explain the absence of QTLs detected in October 2018. Indeed, sample size is known to influence the power of a study to detect QTLs (Hong and Park 2012), and is regularly discussed as one of the most important concerns when designing a mapping experiment (Ashton, Ritchie et al. 2017). Interestingly, two traits (Peduncle Length and Estimated Weight) shared the same QTLs on two linkage groups in the first measurement period (November 2017). This is consistent with the level of high genetic and phenotypic correlations reported for body length and body weight (Valenza-Troubat, Montanari et al. 2021). These findings suggest that selection applied on easily measurable traits such as length will result in the concomitant improvement of more difficult to assess, yet valuable, growth traits (such as weight) in the breeding of trevally.

GWAS identified 113 associations with growth, corresponding to 93 different SNPs spanning 22 LGs, further supporting the hypothesis of a polygenic basis of growth-related traits in trevally. Growth is considered as a complex trait and has been found to be polygenic across the tree of life, in very diverse taxa from plants, like in the model species *Arabidopsis thaliana* (Wieters, Steige et al. 2021), to vertebrates like humans (Sinnott-Armstrong, Tanigawa et al. 2021) and other fish species (e.g. Liu, Sun et al. 2014, Yang, Wu et al. 2020, Debes, Piavchenko et al. 2021). In our study, a lower number of loci were found with the QTL mapping experiment than with the GWAS, which can be explained by the smaller amount of genetic variation represented in the single F_1_ family compared with the overall breeding population, which was derived from thirteen parental individuals. Four QTLs and 15 SNPs were found to have a significant association with more than one trait. This was expected, as the different phenotypes were all deduced from the Peduncle Length and they all then measured the same process underlying growth.

For each trait, the markers found in association were different from a year to another, both in QTL mapping and in GWAS. These differences could be due either to significant loci changing over time, by switching on or off, or to environmental variation, as it was observed in other QTL mapping studies that investigated traits highly affected by the environment (Sun, Niu et al. 2017, Ashton, Ritchie et al. 2019). However, there were some hot spot regions of 0.5 Mb that contained SNPs associated to different traits and different years. Noteworthy is the chromosomic region at the top of LG5, where two hot spots were found, one between 0 and 0.5 Mb (for ΔEW and PL in 2018, and H75 and ES in 2019) and one between 1.4 and 1.8 Mb (for ΔEW, H25, H50 and PL in 2019). Future studies should investigate these further. In addition to these hot spots, we also identified SNP regions that were significant in both QTL and GWAS analyses. Specifically, by comparing the relative physical positions of the SNPs, two regions found with QTL mapping appeared to be in close proximity with two significant SNPs identified by GWAS. In particular, the QTLs for H25, PL, and EW measured in 2017 spanned the 4.45-4.50 Mb region on chromosome 3 (corresponding to 23.20-24.90 cM on LG3) and SNP trevally000114_4956408, located at 4.96 Mb on chromosome 3, was found to be significant for PL in 2019 with GWAS; and the QTLs for ΔH75 and ΔPL in 2019, encompassing the 25.18-25.21 Mb region on chromosome 14 (17.02-17.59 on LG14), are close to SNP trevally001025_25939455, located at 25.94 Mb on chromosome 14 and significant for ΔH50 and ΔPL in 2019 (Figure 5, Table 3 and Table S1). Being identified with two different statistical analyses and for multiple traits across two years, these two regions are then of particular interest for the understanding of the genetic determinism of growth in trevally. A BLAST search of the 100 kb regions flanking those markers against the NCBI nucleotide database did not return any sequence similarities, indicating that they are located in non-coding or in non-annotated regions. Intergenic regions can still have a functional role in gene expression and regulation (Wyrick and Young 2002). For example, these SNPs could be located in an intron that acts as a regulatory region (promoter, enhancer, silencer or insulator).

**Figure 5:**
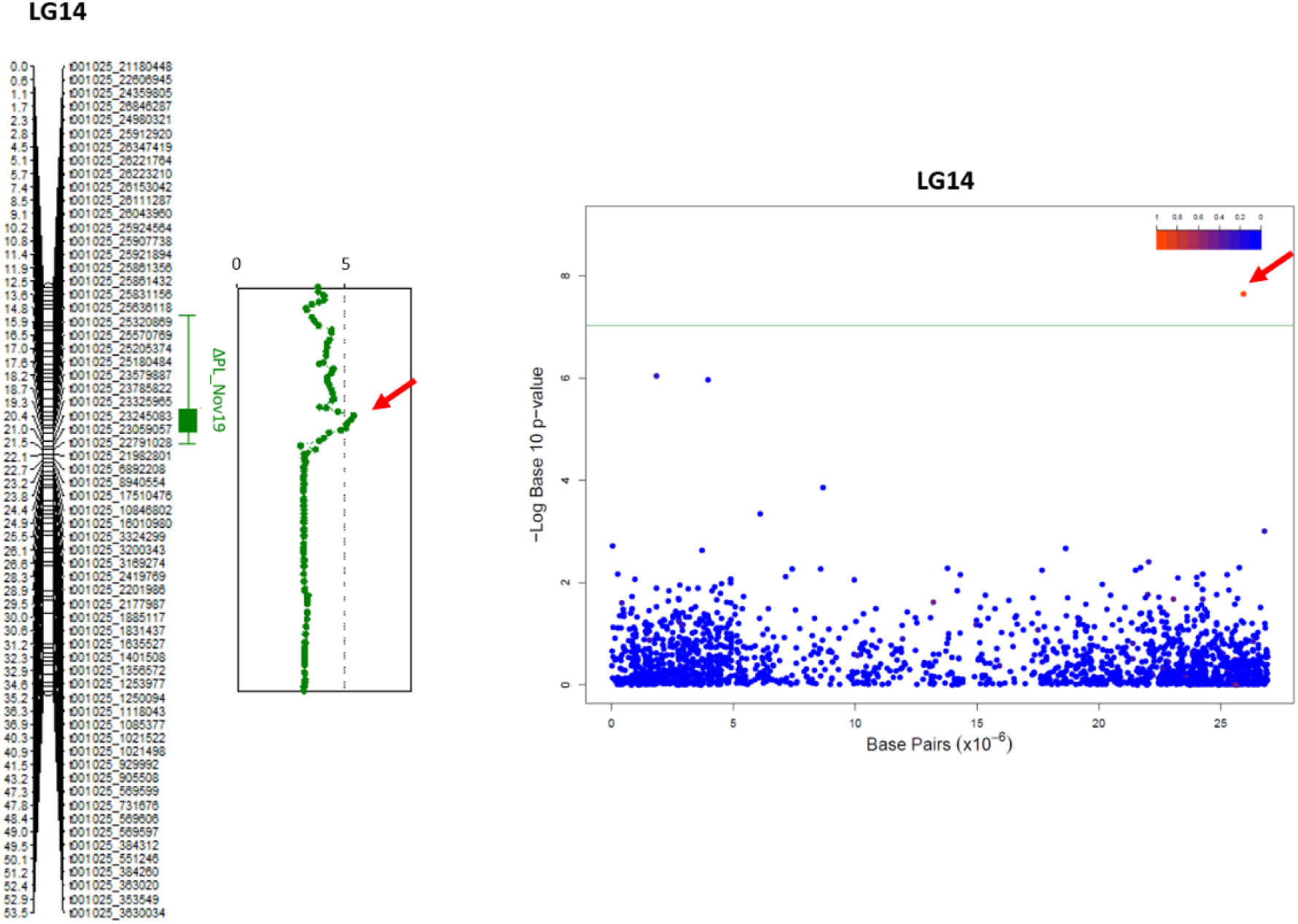
Comparison of Quantitative Trait Loci (left) and SNP associated (right) with the net gain in peduncle length (ΔPL) in November 2019.

Compared with the heritability estimates found in Valenza-Troubat et al. (2021), the present analysis still seems underpowered. The range of heritability estimates was moderate to high (0.67 ± 0.05 to 0.76 ± 0.06) for the measured traits (H25, H50, H75, PL and EW) and moderate (ranging from 0.28 ± 0.07 to 0.68 ± 0.07) for the net gain traits (ΔH25, ΔH50, ΔH75, ΔPL and ΔEW), and it remained consistent throughout the experiment. In the present study, PVEs ranged from 22.1% to 47.36% for the QTL mapping study and from 4.74% to 46.81% for GWAS, indicating that neither techniques were able to capture all of the genetic components of growth traits (Table 4). Additional genetic interactions, other than additive effects, should be considered. While heritability is a key feature of a trait indicating its potential for improvement via selection, polygenic traits are often influenced by non-additive genetic effects such as dominance or epistasis (Glover, Solberg et al. 2017). Understanding the genetic mechanisms that underlie a trait is an important part of explaining phenotypic diversity. This is particularly relevant when looking at traits related to fitness (e.g. growth, shyness, foraging or predator awareness) in populations that are undergoing domestication but still occasionally interbreed with wild conspecifics (e.g. when new broodstock is caught and added).

**Table 4:** Comparison of heritability estimates found in Valenza-Troubat et al. (2021) and Percentage of Variance Explained (PVE) by Quantitative Trait Loci (QTL) found in interval mapping and markers from the Genome Wide Association Study (GWAS). Data are shown for phenotypes height at 25%, 50% and 75% of the body (H25, H50 and H75 respectively), Peduncle Length (PL), estimated weight (EW) and net gains traits associated (ΔH25, ΔH50, ΔH74, ΔPL and ΔEW respectively) in November 2017 (Nov17), October 2018 (Oct18) and November 2019 (Nov19).

| Time | Trait | Heritability | PVE QTL (%) | PVE GWAS (%) |
| --- | --- | --- | --- | --- |
| Nov17 | H25 | 0.67±0.05 | 22.1 | 18.4 |
| Nov17 | H50 | 0.75±0.05 | -- | 24.2 |
| Nov17 | H75 | 0.73±0.05 | 25.3 | 38.5 |
| Nov17 | PL | 0.74±0.05 | 45.3 | 28.8 |
| Nov17 | EW | 0.75±0.05 | 47.4 | 42.3 |
| Oct18 | H25 | 0.74±0.05 | -- | 38.6 |
| Oct18 | H50 | 0.75±0.05 | -- | 24.2 |
| Oct18 | H75 | 0.68±0.05 | -- | 4.7 |
| Oct18 | PL | 0.70±0.05 | -- | 40.6 |
| Oct18 | EW | 0.70±0.05 | -- | 5.3 |
| Oct18 | ΔH25 | 0.46±0.05 | -- | 10.9 |
| Oct18 | ΔH50 | 0.52±0.05 | -- | 43.5 |
| Oct18 | ΔH75 | 0.28±0.05 | -- | -- |
| Oct18 | ΔPL | 0.47±0.05 | -- | 46.8 |
| Oct18 | ΔEW | 0.63±0.05 | -- | 18.0 |
| Nov19 | H25 | 0.73±0.05 | -- | 20.3 |
| Nov19 | H50 | 0.72±0.05 | -- | 41.6 |
| Nov19 | H75 | 0.68±0.05 | -- | 31.2 |
| Nov19 | PL | 0.69±0.05 | -- | 26.4 |
| Nov19 | EW | 0.72±0.05 | -- | 21.4 |
| Nov19 | ΔH25 | 0.56±0.05 | -- | 16.1 |
| Nov19 | ΔH50 | 0.54±0.05 | -- | 15.2 |
| Nov19 | ΔH75 | 0.43±0.05 | 39.7 | -- |
| Nov19 | ΔPL | 0.51±0.05 | 31.0 | 21.7 |
| Nov19 | ΔEW | 0.68±0.05 | -- | 11.9 |

### Future directions

In this study, the combination of QTL mapping and GWAS enabled the identification of genomic regions that control growth in a large captive trevally breeding population. From a farming perspective, parameters such as stocking density (Irwin, O’halloran et al. 1999) or feed availability and quality (Holm, Refstie et al. 1990) can be carefully managed to accelerate fish growth in a land-based aquaculture facility. However, when aiming to develop new species for aquaculture, it is fundamental to understand the genetic architecture of commercially important traits that can potentially be enhanced through selective breeding. The findings of this study provide a useful framework for determining the genetic basis for growth traits in trevally. The identification of multiple QTLs through QTL mapping and genetic markers commonly involved in growth-related traits via GWAS, represents an essential step for the establishment of marker-assisted selection (MAS) breeding programme for trevally. Fine mapping, confirmation and annotation of relevant regions will bring a deeper understanding of the genetic architecture of growth.

## Data Availability Statement

Table S1 contains a list of SNP-trait associations detected with GWAS for height at 25%, 50% and 75% of the peduncle length (H25, H50 and H75 respectively), Peduncle Length (PL), Estimated Weight (EW) and net gain traits (ΔH25, ΔH50, ΔH75, ΔPL, ΔEW) in November 2017 (Nov17), October 2018 (Oct18) and November 2019 (Nov19). For each SNP the table show the physical position on the linkage group (LG), the p-value, the minor allele frequency (MAF), the False Discovery Rate (FDR) adjusted p-value and the percentage of variance explained (PVE).

Trevally (araara) are a taonga (treasured) species to Māori, the Indigenous people of Aotearoa New Zealand. All genomic data obtained from taonga species have whakapapa (genealogy that includes people, plants and animals, mountains, rivers and winds) and are therefore taonga in their own right. These data are tapu (sacred) and tikanga (customary practices, protocols, and ethics) determine how people interact with these data. Thus, all the genomic data have been deposited in a managed repository that controls access. Raw and analyzed data are available through the Genomics Aotearoa data repository at https://repo.data.nesi.org.nz/. This was done to recognise Māori as important partners in science and innovation and as inter-generational guardians of significant natural resources and indigenous knowledge.

## Acknowledgements

We would like to acknowledge the PFR staff that assisted with the breeding and husbandry operation for the trevally populations; in particular Warren Fantham who oversees the larvae rearing of finfish and Therese Wells who manages the post-juvenile husbandry. This research was funded through the MBIE Endeavour Programme “Accelerated breeding for enhanced seafood production” (#C11X1603) to MW.

